# The distribution of fitness effects among synonymous mutations in a gene under selection

**DOI:** 10.1101/553610

**Authors:** E. Lebeuf-Taylor, N. McCloskey, S.F. Bailey, A. Hinz, R. Kassen

## Abstract

The fitness effects of synonymous mutations, nucleotide changes that do not alter the encoded amino acid, have often been assumed to be neutral, but a growing body of evidence suggests otherwise. We used site-directed mutagenesis coupled with direct measures of competitive fitness to estimate the distribution of fitness effects among synonymous mutations for a gene under selection. Synonymous mutations had highly variable fitness effects, both deleterious and beneficial, resembling those of nonsynonymous mutations in the same gene. This variation in fitness was underlain by changes in transcription linked to the creation of internal promoter sites. A positive correlation between fitness and the presence of synonymous substitutions across a phylogeny of related Pseudomonads suggests these mutations may be common in nature. Taken together, our results provide the most compelling evidence to date that synonymous mutations with non-neutral fitness effects may in fact be commonplace.

## Main Text

Our ability to use DNA sequence data to make inferences about the evolutionary process from genes or genomes often relies on the assumption that synonymous mutations, those that do not result in an amino acid change, are neutral with respect to fitness. Yet there is compelling evidence that this assumption is sometimes wrong: comparative (1) and experimental (2) data show that synonymous mutations can have a range of fitness effects from negative to positive, and can even contribute to adaptation (3–6). A range of mechanisms including codon usage bias, altered mRNA structure, and the creation of promoter sequences could lead to changes in the rate or efficiency of transcription, translation, and/or protein folding and/or expression that, in turn, impact fitness (7). The specific mechanism notwithstanding, it is clear that synonymous mutations are not always neutral; the degree of variability in their fitness effects, and how often they contribute to adaptation, remains unknown.

As a first step towards answering these questions, we estimated the distribution of fitness effects (DFE) for 39 synonymous, 65 nonsynonymous, and 6 nonsense substitutions at 34 sites along *gtsB*, a gene that codes for a membrane-bound permease subunit of an ABC glucose-transporter in the Gram-negative bacterium *Pseudomonas fluorescens* SBW25. Single nucleotide mutants were generated through site-directed mutagenesis and competed against the ancestor strain in glucose-limited medium. We previously reported on two beneficial synonymous mutations in this gene recovered from a population that had evolved for ~1000 generations in glucose-limited medium and confirmed that *gtsB* is a target of selection under these conditions (3). Visual inspection of the DFEs for nonsynonymous and synonymous mutations (Fig 1A) reveals they are similar, with both having modes close to neutrality (*w* ~1) and substantial variation that includes mutants with both positive and deleterious effects. However, the distributions differ significantly (*P* = 0.0002 based on a bootstrapped estimate of the Kolmogorov-Smirnov D-statistic from 10,000 permutations) due to the presence of a handful of strongly deleterious nonsense mutations in the non-synonymous set that presumably a produce truncated, non-functional protein. Remarkably, the DFEs for beneficial nonsynonymous and synonymous mutations are indistinguishable (bootstrapped K-S test, P = 0.59), suggesting that both kinds of mutation could contribute to adaptation. The combined DFE for both kinds of beneficial mutations is approximately L-shaped, with many mutations of small effect and a few of large effect (Fig 1B), as expected from theory (8–10). More formally, the DFE among beneficial mutations is significantly different from an exponential distribution (likelihood ratio test, P = 0.0077) and falls within the Weibull domain of the Generalised Pareto Distribution (K = −0.37), suggesting the existence of a local fitness optimum similar to what has been seen previously for nonsynonymous mutations (11,12).

**Fig. 1.**
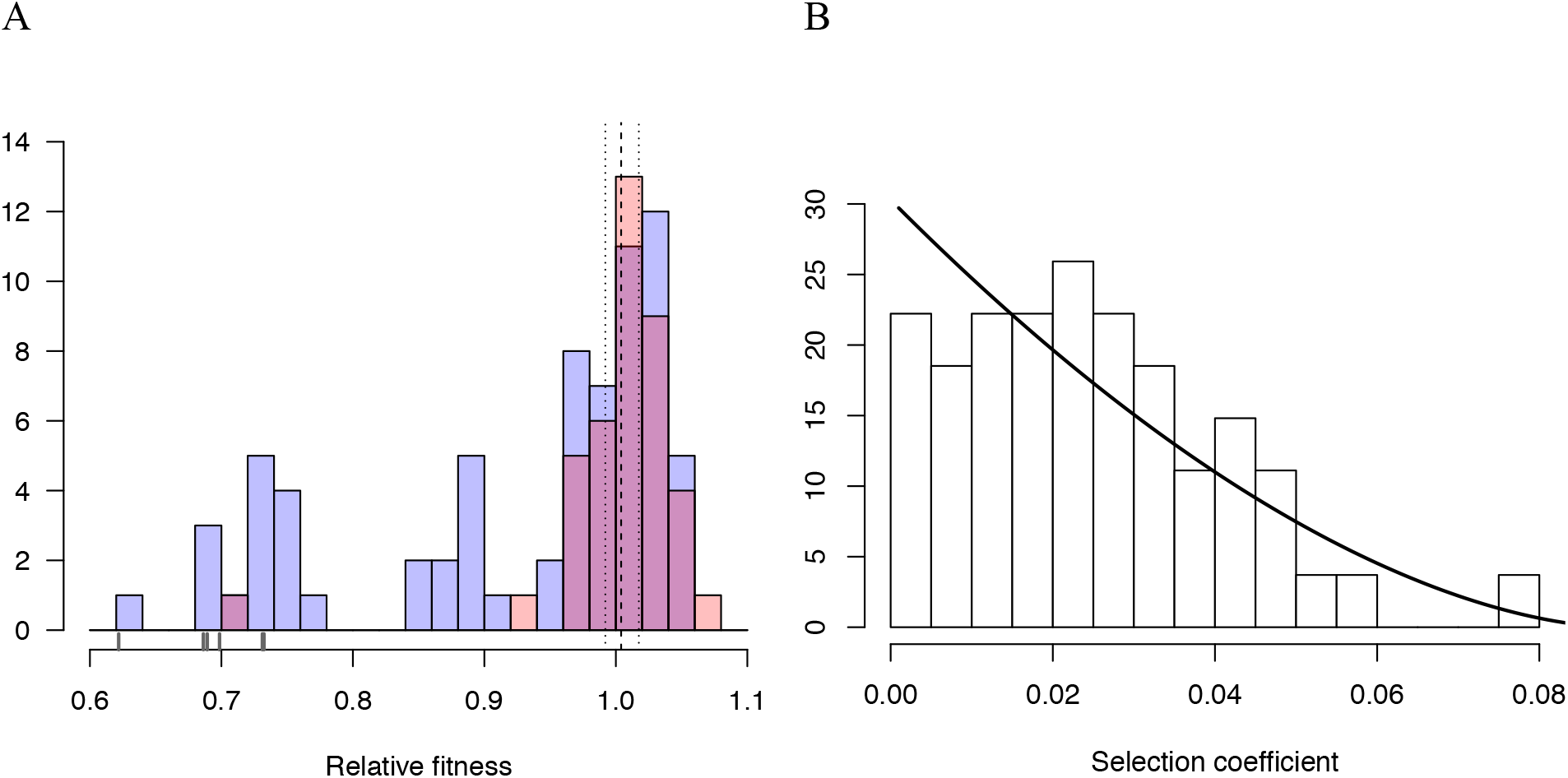
Distributions of relative fitness effects of *gtsB* points mutations in low glucose media. (A) Counts of nonsynonymous (blue; n = 71) and synonymous (red; n = 39) mutations display a wide range of fitness effects, with ticks under the bars indicating the relative fitness values of nonsense mutations. Dashed and dotted lines show the mean relative fitness of the wild-type (WT) competed against the marked competitor. (B) The DFE of beneficial-effect mutations (proportions; pooled synonymous and nonsynonymous samples, n = 55) is fit by a k value of − 0.35, which corresponds to the Weibull domain of attraction of the Generalised Pareto Distribution. On this normalized histogram (total area = 1), relative fitness values are shifted to the smallest observed value and expressed as selection coefficients.

What cellular processes underlie the wide range of fitness effects observed here? Our fitness data allow us to test some of the leading hypotheses through *in silico* analyses. Fitness could be higher if synonymous mutations result in codon usage that is more closely aligned with that of highly expressed genes. Alternatively, it has been suggested that suboptimal codon usage within the first ~50 codons -- the translational ramp -- is required to ensure efficient translation initiation (13,14), suggesting that higher fitness should be associated with the introduction of rarer codons close to the start of a gene. We could find no evidence for either explanation in our sample: a regression of fitness on distance from the start codon of *gtsB* was not statistically significant (permutation of residuals, P = 0.20) nor was the change in codon adaptation index (CAI; P = 0. 40) or tRNA adaptation index (tAI; P = 0.53), both measures of the degree of codon usage bias. In fact, we find no evidence of a translational ramp in the WT *gtsB* (adj. R^2^ = 0.0058, P = 0.21); further, the interaction of codon position with CAI or tAI does not yield significant results (P = 0.45 and 0.90, respectively).

It has also been suggested that synonymous mutations could impact fitness through their effects on mRNA transcript secondary structure and hence the rate and fidelity of translation. Higher fitness could result from faster translation due to transcripts that are less thermodynamically stable and, so, more accessible to the ribosome during translation (15) or from more efficient translation due to more stable mRNAs that persist longer due to slower degradation rates (16). A linear model linking change in mRNA stability and fitness is significant for the nonsynonymous subset of mutations (permutation of residuals, P = 0.0039), although the effect is weak (adj. R^2^ = 0.11) and largely driven by less stable, highly deleterious mutations. We could not detect a relationship between change in mRNA stability and fitness for synonymous mutations, even when we account for the possibility of strong 5’ end secondary structures by adding a position term reflecting distance from the start codon (17).

The absence of any relationship between synonymous mutation fitness and codon usage bias or mRNA stability, both measures affecting translation, suggests that fitness effects stem from changes in transcription. Testing this hypothesis requires comparing estimates of transcript and protein abundance, the difference being a measure of the effect of translation. We evaluated mRNA and GtsB protein levels by proxy via the insertion of a yellow fluorescent protein (YFP) bioreporter into the WT or mutant *gtsB* background just before or just after the stop codon. The former construct (a translational fusion) produces a single transcript where *gtsB* and YFP are translated together; the latter, with YFP inserted after the *gtsB* stop codon (a transcriptional fusion), results in *gtsB* and YFP translated separately. A mutation upstream of these fusions that leads to increased translation, but not transcription, is expected to generate a higher YFP expression level in the translational fusion compared to the transcriptional fusion. The expression levels of these different constructs relative to the WT are shown in Figure 2B for the two synonymous mutations (A15A, G38G) and a third, independently evolved non-synonymous mutation (A10T) recovered from the original experiment by Bailey et al (2014). Transcription is elevated in all three mutants relative to the WT but we could not detect any additional effect of translation in the two synonymous mutations, although there is a modest increase in expression associated with translation for the nonsynonymous mutation. These results suggest that the primary effect of these synonymous mutations is on levels of transcription rather than translation.

**Fig. 2.**
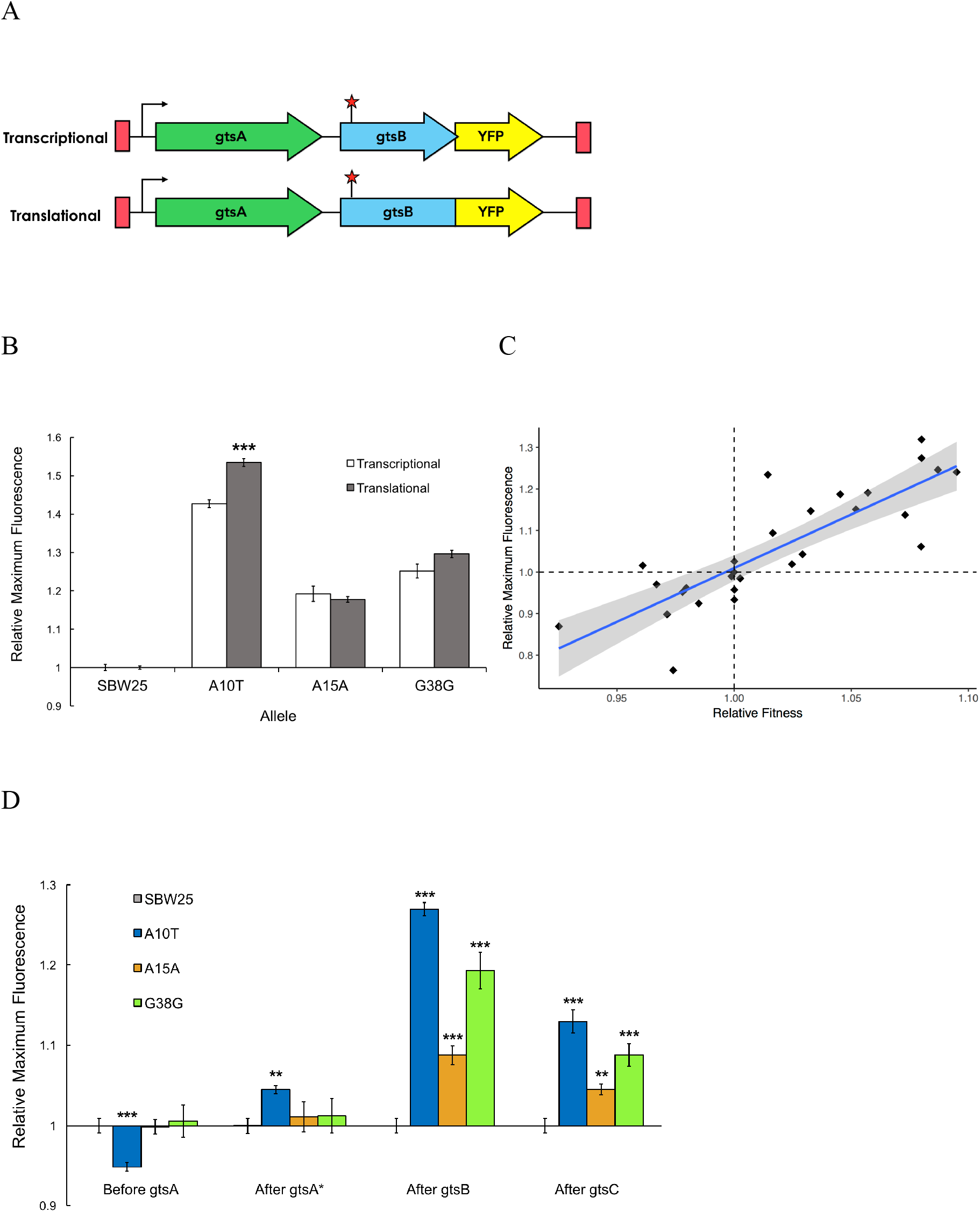
Comparison of transcriptional and translational effects of the evolved mutants and correlation with relative fitness. (A) Schematic showing the sites of YFP insertion for transcriptional and translational fusions. (B) Comparison of maximum YFP expression (± SEM) from transcriptional and translational YFP fusions at the Tn7 site for the WT (n = 14 replicates) and evolved mutants (n = 7, 7, and 6 technical replicates, respectively). Significance with respect to transcriptional fusion: *** P < 0.001. (C) Linear regression of fluorescent signal of YFP transcriptional fusions as a proxy for transcript levels and relative fitness measures for a subset of synonymous mutations (n = 27). Grey shading indicates the 95% confidence interval for the regression (adjusted R^2^ = 0.69, P < 0.001). (D) Protein expression across the *gts* operon for evolved mutants. Maximum fluorescence (± SEM) of the YFP transcriptional fusions at different loci in the *gts* operon relative to SBW25. ** P < 0.01, *** P < 0.001.

Two additional lines of evidence point to changes in transcription levels as the likely proximate cause of variation in fitness among our synonymous mutations. First, there is a strong positive relationship between transcript abundance and relative fitness for 27 synonymous mutations (including the A15A and G38G mutations examined above) (Fig 2C; R^2^ = 0.691, P = 1.46 × 10^−8^). Notably, the range of the regression includes mutants with both negative and positive fitness effects, suggesting that the link between transcript abundance and fitness is not limited to beneficial synonymous mutations alone. Second, Figure 2D shows that the increased transcription caused by A15A, G38G, and A10T extends downstream to *gtsC* (P < 5.0 × 10^−5^) but not upstream to *gtsA*, which remains largely unaffected (P > 0.60). These synonymous mutations thus have polar effects on transcription that extend beyond the gene in which they occur. Taken together with our previous observation that overexpression of WT *gtsB* increases fitness only when the rest of the *gts* operon is also overexpressed (3), these results suggest that co-expression of downstream genes is necessary for increased fitness in this system.

What mechanism accounts for the observed changes in transcription and fitness among the synonymous mutations? Previous work has shown that synonymous mutations can generate beneficial effects by creating novel promoters in regions upstream of a gene under selection (18,19). At face value, this mechanism cannot explain our results since we observe a range of fitness effects for synonymous mutations along the entire length of *gtsB*. However, the existence of polar effects on transcription suggests that some synonymous mutations in *gtsB* might be playing a similar role by creating internal promoters causing changes in expression of the downstream genes *gtsC* or *gtsD*. To evaluate this idea, we used Softberry BPROM online software to search the entire *gts* operon for internal sigma 70 bacterial promoter sequences in the ancestral sequence and mutant sequences. We find relatively few hits in our collection, perhaps because BPROM searches for promoters using *Escherichia coli* rather than *P. fluorescens* consensus sequences; however, among the top five hits is a predicted promoter sequence spanning codons 30-42 that includes G38G and 39-3T, the latter being the synonymous mutation with highest fitness in our collection. Both mutations, and an additional beneficial synonymous mutation at 232-3T, result in predicted −10 promoter sequences that are more closely aligned to the −10 consensus sequence for *P. aeruginosa* (TATAAT) than the WT. Notably, there is a tendency for promoter strength to vary positively with both transcription and fitness (Fig 3), although this effect is based on just six mutations and is not significant (permutation of residuals, P = 0.17 and 0.11, respectively). These results suggest that fitness changes associated with these synonymous mutations could be caused by the ability of transcription factors to bind to promoter-like sequences in *gtsB* and alter transcription of downstream genes.

**Fig. 3.**
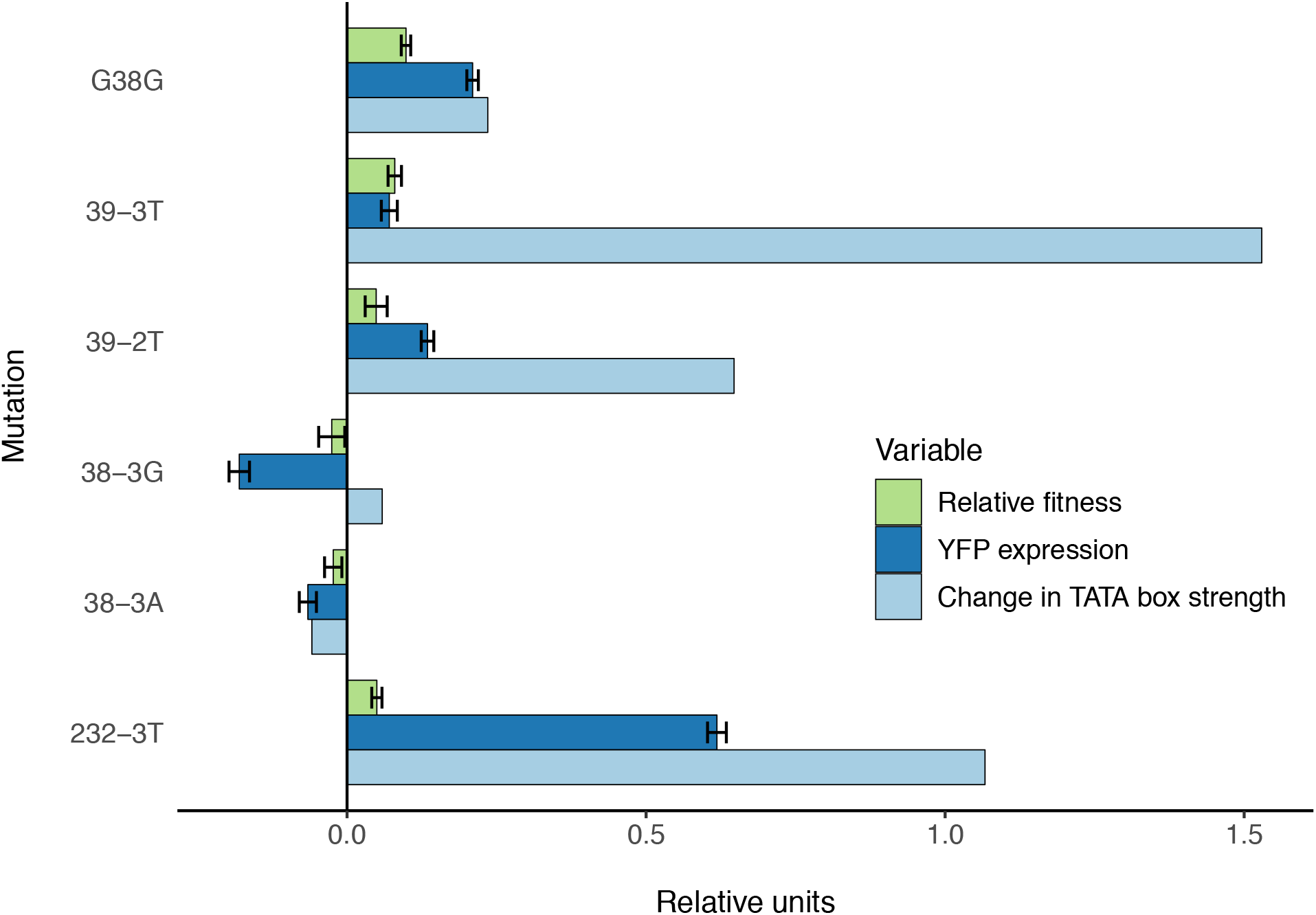
Potential mechanism for fitness differences at different loci in *gtsB*. Bars represent the mean of each variable in units relative to the WT. Experimental relative fitness and transcriptional expression of YFP measures include standard error.

How often do these synonymous mutations contribute to adaptation in more natural settings beyond the highly contrived conditions we have studied here? We can get partway towards an answer by asking whether fitness *in vitro* predicts prevalence of a mutation across a phylogeny of pseudomonads. We generated a phylogeny of 77 strains closely related to SBW25 and converted the probability of observing a given mutation to a binary variable based on its presence or absence in the phylogeny. For our entire synonymous and nonsynonymous sample, we find a positive relationship between the presence of a particular mutation in the phylogeny and its fitness in glucose-limited medium (Fig 4, P = 0.0210). Notably, our highest fitness mutation, 39-3T, which is synonymous, arises independently multiple times across the phylogeny. These results lend support to the idea the variation in fitness effects observed here are not an idiosyncratic result of life in a laboratory environment. Rather, the synonymous mutations conferring the highest fitness effects may often contribute to adaptation in more complex, and more natural, environments as well.

**Fig. 4.**
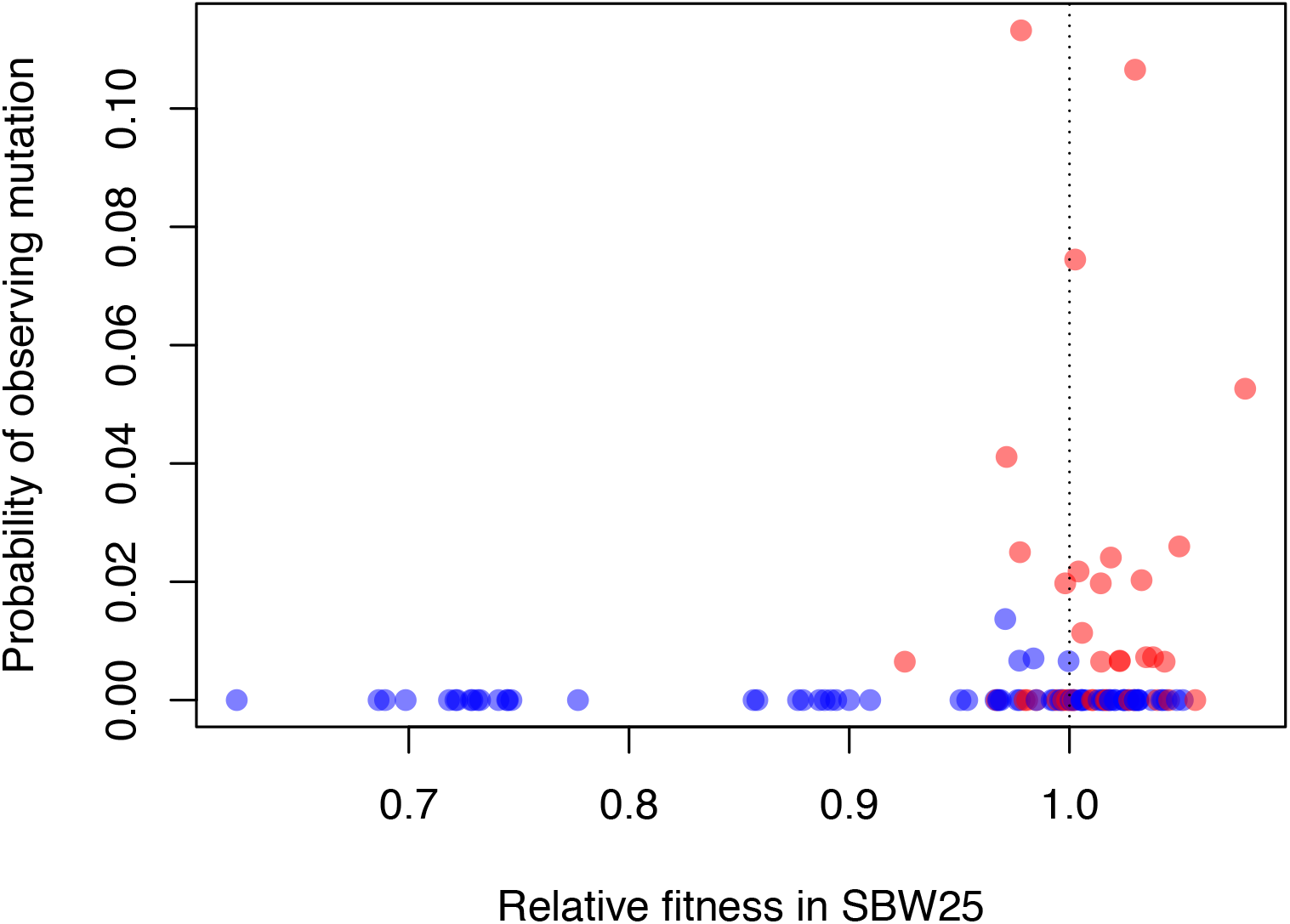
Beneficial synonymous mutations (red) are often observed in the phylogeny of related pseudomonads, while nonsynonymous mutations are less so (blue). There was a significant logarithmic relationship between the probability of observing a given mutation as a binary variable (present/absent) and relative fitness (P = 0.0121).

We have shown that, contrary to what has often been assumed, the DFE among synonymous mutations can be highly variable and include both deleterious and beneficial mutations. In fact, aside from the absence of strongly deleterious mutations associated with premature stop codons, the DFE of synonymous mutations is strikingly similar to that of nonsynonymous mutations and is formally indistinguishable from it if we consider only beneficial mutations. Taken together with the observation of a positive correlation between *in vitro* estimates of fitness of a given mutation and its prevalence among sequenced isolates, these results suggest that these synonymous mutations can, and sometimes do, contribute to adaptation.

The cause of the fitness variation among synonymous mutations observed here stems from changes to transcription that impact downstream genes in the same operon. Whether these transcriptional effects occur by changing an internal promoter sequence, as our data suggests, or through some other, still undiscovered mechanisms, remains to be elucidated. It is notable that promoter-associated effects on transcription have been shown to underlie the fitness effects of synonymous mutations in two other microbial systems (18,19), suggesting this mechanism may be quite general, at least for organisms with operon-like genetic architectures. Nevertheless, others have pointed to changes in translational efficiency associated with the accessibility of mRNA near a start codon as the primary mediator of fitness in *Salmonella enterica* (5) and synonymous mutations are known to impact fitness in a wide range of organisms beyond prokaryotes (1,20,21). Uncovering the full spectrum of mechanisms by which synonymous mutations impact fitness, and how often they contribute to adaptation, remains a major task for the future.

## Materials and Methods

### Culture conditions

*E. coli* was grown on Luria-Bertani (LB), X-gal sucrose, or tetracycline media. *Pseudomonas fluorescens* SBW25, which was used as the ancestral strain, was grown on LB or X-gal minimal salts media (48mM Na_2_HPO_4_, 22mM KH_2_PO_4_, 9mM NaCl, 19mM NH_4_Cl, 2mM MgSO_4_, 0.1mM CaCl_2_) with glucose (53 mM), succinate (80 mM) or mannitol (53 mM) as indicated. Media were supplemented with 5-bromo-4-chloro-3-indolyl-b-D-galactopyranoside (X-gal) at 40 mg/ml. Antibiotics were used at the following concentrations: 100 mg/ml nitrofurantoin (Nf), 100 mg/ml ampicillin (Ap), 10 mg/ml tetracycline (Tc).

### Construction of *gtsB* mutagenesis vector

The *P. fluorescens* allelic exchange vector pAH79 (3) was modified for rapid generation of mutant *gtsB* alleles by Golden Gate assembly (GGA) of polymerase chain reaction (PCR) amplicons. A three-part ligation between a digested pAH79 derivative (including BglII and SpeI sites as well as a BsaI cloning site compatible with that of oligo F2-gtsB-F), *lacZα* amplicon (BglII and MfeI), and SBW25 amplicon spanning 114 to 865 bp downstream of the *gtsB* stop codon (MfeI and XbaI) yielded the final *gtsB* mutagenesis vector.

### Site-directed mutagenesis of *gtsB*

Site-directed mutagenesis of *gtsB* (*PFLU4845*) was accomplished by cloning *gtsB* alleles with a single mutation into a vector as described above, generating an *E. coli* library, followed by allelic replacements in SBW25. Primers were designed to introduce mutations at 112 sites spanning the *gtsB* gene. Sequences from 715 base pairs upstream to 173 base pairs downstream of *gtsB* were amplified as two PCR fragments, one of which contained a threefold degenerate polymorphism introduced by a mutagenic primer. BsaI recognition sequences were included in each primer to enable seamless ligation between the PCR products and mutagenesis vectors using GGA (22). Cloning reactions were transformed into *E. coli* DH5α λpir following the Inoue method for chemical transformation (23) with selection on ampicillin.

Transformations yielded libraries of *E. coli* strains for introduction of mutations into SBW25. Recombination of each mutant *gtsB* allele into the chromosome was selected for in two steps: selection for Tc^R^ followed by selection for sucrose resistance as previously described (3). We used an SBW25 recipient strain in which the native *gtsB* was replaced by *lacZ*, allowing us to use blue-white screening on LB 5% sucrose X-gal agar to identity recombinants in which *lacZ* was replaced by the vector-encoded mutant *gtsB*. The sucrose-resistant white colonies were used as PCR templates for amplification of the *gtsB* locus using an M13F-tagged primer (Table 1), for sequencing by the McGill University and Genome Quebec Innovation Centre.

**Table 1.**
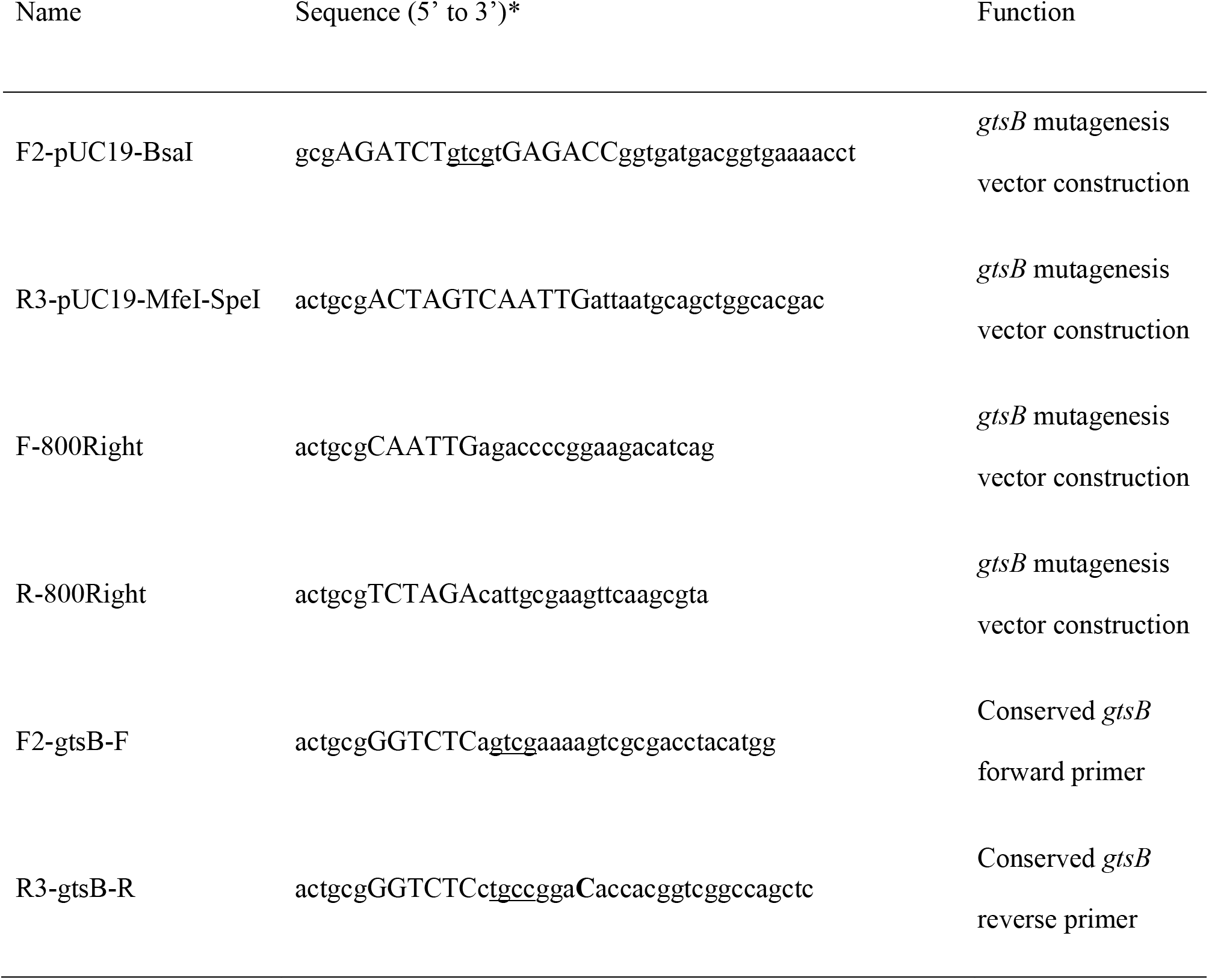
Oligonucleotides used in this study. Restriction enzyme recognition sequences are capitalized. BsaI overhangs are underlined. Introduced mutations are in bolded capital letters.

### DNA extraction and sequencing

Strains were grown in LB liquid media overnight; genomic DNA was extracted using the QIAGEN DNeasy Blood & Tissue Kit. Sequence data were generated on the Illumina MiSeq platform with paired-end reads using the Nextera XT kit. Reads generated were approximately 300 bp in length.

### Reference-based mapping and variant calling

The genome of *P. fluorescens* SBW25 is publicly available from NCBI (PRJEA31229). A modified version of the bioinformatics pipeline described in Dettman et al. (2012) (24) was used to analyse the reads. Briefly, reads were trimmed using Popoolation (ver. 1.2.2, (25)) with a Phred quality threshold of 20 and a minimum retention length of 75% of original read length. Trimmed reads were then mapped to the SBW25 reference genome using Novoalign (ver. 3.02.08, www.novocraft.com). Single nucleotide polymorphisms and indels were annotated using Samtools (ver. 1.9, (26)), BCFtools (ver. 1.9), VarScan (ver. 2.3.7, (27)), and snpEff (ver. 4.0, (28)). Read and alignment quality were assessed using FastQC (ver. 0.11.7, www.bioinformatics.babraham.ac.uk/projects/fastqc/). Sequence data are available from the NCBI Short Read Archive under BioProject PRJNA515918: Pseudomonas fluorescens SBW25 gtsB mutants.

### Competitions

Competitions were performed as outlined in Lenski *et al* (29) on four to six replicates (genetically identical clones) of 110 mutant strains. These replicates provide a measure of the variability inherent in our experimental procedure, and as thus considered technical replicates. All strains, including SBW25-*lacZ*, were removed from storage at −80°C and grown overnight at 28°C on LB agar. Single colonies were inoculated into 2 mL LB broth for overnight incubation at 28°C under shaking conditions. Each mutant strain was transferred into minimal glucose media for a 24h acclimation period at 28°C, then mixed in a 1:1 volumetric ratio with SBW25-*lacZ* and inoculated into 2 mL of minimal media with glucose. Initial and final aliquots from mixed cultures were frozen in 20% glycerol after 1 and 24 hours’ growth and plated on minimal agar with glucose. Only plates containing 30 or more colonies of each strain were included. Relative fitness was calculated using *w* = (*f_final_*/*f_initial_*)^(1/doublings), where *f_final_* and *f_initial_* are ratios of the frequency of mutant to the frequency of SBW25-*lacZ* strain after and before the competition. The number of doublings was estimated from the dilution factor and corresponded to ~ 6.7 or 13.2 generations depending on the dilution factor. The effect of the *lacZ* marker was tested by competing SBW25-*lacZ* against the WT with each batch of competitions. The mean relative fitness of SBW25-*lacZ* was 1.005 ± 0.0007 SEM.

### Estimating codon preference and mRNA stability

In order to estimate the change in codon bias attributable to each synonymous mutation, we compared the CAI value of the mutant to the WT using SBW25 ribosomal protein genes as a reference (30). The ‘cai’ function in the ‘seqinr’ package in R (31) was used to calculate change in CAI at each site. tAI values were calculated by inputting tRNA gene copy number, a proxy for tRNA expression (13), in the stAI_calc_ interface (32). As per previous work (15), we predicted the most likely folding energy of 42-nucleotide windows centred on each mutation using the ‘mfold’ server (33).

### Comparison and characterisation of DFEs

All statistical analyses were conducted in R Studio (version 1.0.136; www.rstudio.com). Six nonsense mutations were omitted from our analysis, since they likely do not result in a complete protein. We compared synonymous and nonsynonymous DFEs for all mutations and for the subset of beneficial mutations by bootstrapping the Kolmogorov-Smirnov (K-S) statistic. We found no significant difference between K-S values for nonsynonymous and synonymous beneficial mutations, so we pooled the data to infer the properties of the tail distribution following the method outlined in Beisel *et al* (35). Relative fitness values were transformed to selection coefficients by subtracting 1; we shifted the threshold to the smallest observed selection coefficient, as suggested by Beisel *et al*. Using the ‘GenSA’ package in R, we estimated the optimal value of the scale parameter *τ*, which characterises the stretch of the distribution, with *κ* (the tail parameter) set to 0, corresponding to an exponential distribution; in the alternative model, optimal *τ* and *κ* values were calculated without restricting *κ*. A likelihood ratio test was used to determine whether the model with the unconstrained *κ* value was a better fit than the exponential distribution.

### Statistical analysis

Permutation of residuals was used as per Still and White (1981) (34) to test for significant relationships between explanatory variables and fitness. We tested for significant differences in YFP expression between mutant and WT alleles using a two tailed T-test assuming equal variance. Threshold for significance was α=0.05.

### Transcriptional and translational fusion of YFP at Tn7 site

Transcriptional and translational fusion constructs were generated using GGA (22) and the use of site-specific mini-Tn7 transposon (36,37). A 2.6 kb PCR product was amplified from template DNA containing the target locus, and included the 346 bp promoter region of *gtsA*, the open reading frame of *gtsA* and the open reading frame of *gtsB*. This PCR product and the downstream YFP fusion PCR product were seamlessly ligated together into the Tn7 vector (mini-Tn7T-Gm) through GGA.

The YFP transcriptional fusion after *gtsB* required additional modifications due to the eight nucleotide overlap between the stop codon of *gtsB* and start codon of *gtsC*. To preserve the *gtsB* sequence and predicted *gtsC* ribosomal binding site, the start codon of the YFP transcriptional fusion was inserted in-frame after the first four codons *gtsC*. Translational YFP fusions had a 6-glycine ((GGC)6) linker sequence (38) between the second-last codon (302) of *gtsB* and the second codon of YFP, which removed both the stop codon of *gtsB* and start codon of YFP to create a single peptide. The 3′ UTR of both the transcriptional and translational YFP fusions includes an intrinsic transcriptional terminator. Tn7 vectors were transformed into *E. coli* DH5α λpir following the Inoue method, then incorporated into SBW25 and verified as described in Bailey *et al* (2014).

### Phylogeny construction

A phylogeny was constructed using full DNA sequences of *rpoB, rpoD*, and *gyrB* for 77 closely related Pseudomonas strains obtained from NCBI, as per Gomila *et al* (39). The concatenated sequences were aligned using NCBI’s BLAST. MEGA7 (40) was used to build the tree based on the aligned concatenated sequences using maximum likelihood to generate a bootstrapped consensus tree (*n* = 500). For each site in the *gtsB* alignment, maximum parsimony was used to estimate the ancestral state at all internal nodes in the phylogeny with the function “ancestral.pars” from the phangorn package in R (41). As our phylogeny had polytomies, the ancestral states were estimated for 100 randomly resolved bifurcating phylogenies and the frequency of the inferred ancestral state at each internal node was calculated. For each site in *gtsB*, the number of evolutionary events was calculated by comparing the inferred state at the beginning and end of each branch and counting the number of transitions. A binary model was fit expressing whether mutations of interest were observed in the phylogeny.

## Acknowledgements

Thanks to N. Rodrigue, A. Schick, and A. Wong for feedback and discussion.

## Author contributions

Experiments were conceived and designed collaboratively under the direction of RK; AH, ELT and NM created mutants and genetic constructs; ELT and NM performed competitions; ELT and SFB analysed the data; ELT and RK wrote the manuscript.

## Data and materials availability

Data are available upon request and will be deposited in Dryad upon publication. Genomic data has been deposited into the NCBI Sequence Read Archive as BioProject PRJNA515918.

